# WSB.*APP/PS1* mice develop age-dependent cerebral amyloid angiopathy, cerebrovascular dysfunction, and white matter deficits

**DOI:** 10.1101/2025.10.08.681261

**Authors:** Olivia J. Marola, Asli Uyar, Kelly J. Keezer, Juan Antonio K. Chong Chie, Kevin J. Elk, Abigail E. Cullen, Kierra Eldridge, Scott Persohn, Jonathan Nyandu Kanyinda, Jennifer D. Whitesell, Julie A. Harris, Paul Salama, Ashley E. Walker, Gregory W. Carter, Michael Sasner, Paul R. Territo, Gareth R. Howell, Kristen D. Onos

## Abstract

**INTRODUCTION:** Growing evidence suggests cerebrovascular deficits, including cerebral amyloid angiopathy (CAA), play a key role in Alzheimer’s disease (AD) pathogenesis. However, these facets of AD are not well understood, due in part to the lack of mouse models that develop robust vascular deficits and CAA. Here, we characterize human-relevant cerebrovascular phenotypes in WSB.*APP/PS1* mice with and without humanized *APOE* alleles.

**METHODS:** AD-relevant cerebrovascular phenotypes in WSB, WSB.*APP/PS1,* WSB.*APOE^2/2^APP/PS1,* WSB.*APOE^3/3^APP/PS1,* and/or WSB.*APOE^4/4^APP/PS1* mice were characterized using immunohistochemistry, transcriptomics, positron emission tomography/computed tomography, and *ex-vivo* analyses.

**RESULTS:** WSB.*APP/PS1* mice exhibited age-related plaque deposition and CAA, significant transcriptomic overlap with human AD, myelin deficits, cerebrovascular/metabolic uncoupling, and altered cerebrovascular morphology. Aged WSB vasculature retained vasoreactivity, but exhibited increased stiffness. Compared to *APOE^2^*, *APOE^4^* expression in WSB.*APP/PS1* mice increased CAA and plaque-associated microglial area.

**DISCUSSION:** These data illustrate the utility of the WSB genetic context to model CAA and uncover vascular contributions to AD.

**Highlights:**

- WSB.*APP/PS1* mice developed CAA with age.
- Transcriptomic profiling revealed significant molecular overlap between human AD and WSB.*APP/PS1* brains.
- Transcriptomics and immunofluorescence suggested age-related myelin deficits in WSB.*APP/PS1* brains.
- WSB.*APP/PS1* brains exhibited neurovascular uncoupling, changes in vascular volume and surface area, and increased permeability changes.
- WSB.*APP/PS1* cerebrovasculature was resilient to loss of responsivity with age but exhibited increased vascular stiffness.
- Humanized *APOE* ε*4* alleles significantly increased CAA and parenchymal plaque-associated microglial area in WSB.*APP/PS1* mice.

**Research in Context:**

1. <u>Systematic review:</u> The authors characterized WSB.*APP/PS1* as a unique human-relevant model of Alzheimer’s Disease and explored several facets of cerebrovascular deficits.
2. <u>Interpretation:</u> WSB and/or WSB.*APP/PS1* mice exhibited human-relevant vascular phenotypes, including CAA, neurovascular uncoupling, increased vascular tree volume, vascular stiffness, and resilience to age-related loss of responsivity. Furthermore, the transcriptomic profile of WSB.*APP/PS1* brains significantly overlaps with signatures observed in human AD. WSB.*APP/PS1* brains exhibited myelin deficits with age. Furthermore, humanized *APOE* ε*2,* ε*3,* and ε*4* alleles significantly modified WSB.*APP/PS1* susceptibility to CAA, plaque deposition, and plaque-associated microglial area.
3. <u>Future Directions:</u> The WSB genetic context will be leveraged to identify specific molecular mechanisms associated with cerebrovascular deficits in AD.

## Background

Alzheimer’s disease and related dementias (ADRD) significantly impact quality of life and the ability to provide self-care. Current global estimates of incidence suggest that the 50 million people living with dementia will grow to exceed 152 million by 2050. Age is the greatest risk factor for ADRD, and this risk is modulated by genetic and environmental factors(*1*). AD is classically defined as the presence of specific pathology in the brain: deposition of amyloid, intracellular neurofibrillary tangles of Tau, neuroinflammation, and neurodegeneration. ADRD diagnosis is difficult, as there are currently few reliable biomarkers. Growing evidence suggests the importance of vascular perturbations in driving AD pathology. One such common AD-relevant vascular pathology is cerebral amyloid angiopathy (CAA)—the deposition of amyloid plaques within the walls of the cerebrovasculature. CAA is present in up to 90% of AD patients(*2*), and is associated with endothelial cell dysfunction, hypoperfusion, production of reactive oxygen species (ROS), neurovascular damage, impaired amyloid clearance, and neurovascular uncoupling(*3, 4*). CAA is also one of the most common risk factors for lobar intracerebral hemorrhages and ischemia in the aging population(*5*). Changes in cerebral blood flow, vascular function, and subsequent metabolic dysregulation are detected in early stages of ADRD(*3, 4*), and are thought to occur prior to the development of CAA. However, despite the importance of vascular contributions to ADRD, the underlying causes of CAA are not well understood. It is vital to decipher the underlying mechanisms behind vascular deficits in ADRD to develop therapeutics that aim to improve vascular health and prevent cognitive decline.

Despite previous work focused on understanding vascular deficits in ADRD, including CAA, advances have been hampered by the lack of translationally relevant animal models(*6*). For instance, the widely used transgenic amyloid model mouse models, such as the 5xFAD or *APP/PS1*, phenocopy parenchymal plaque deposition but lack robust CAA, even at older ages (*6*). We posit this is due in part to the manufactured nature of classically used mouse strains–most commonly C57BL/6 (B6)(*6*). Commonly used laboratory strains were generated through random crosses of European and Asian fancy mouse stocks, and have a mixture of genomic contributions from all three *Mus musculus* subspecies(*7*). Recent work from our lab has leveraged wild-derived mouse strains to uncover genetic underpinnings of resilience and susceptibility to AD pathogenesis(*8–10*). For example, WSB/EiJ (WSB) is a wild-derived mouse strain that was first established by wild mice trapped in Maryland that were sent to The Jackson Laboratory in 1986 and then inbred(*9*). The genomes of these mice are specifically of the sub-species *Mus musculus domesticus*, and critically, harbor a unique genetic context that can aid in the understanding of aging and diseases with complex etiologies(*7, 11*).

To investigate the utility of WSB for modeling ADRD, the *APP/PS1* transgenes were previously backcrossed from B6 to WSB for at least 10 generations(*9*). This initial characterization was performed after seven generations, and ADRD phenotypes assessed at 8 months of age(*9*). At this age, in comparison to the commonly used B6.*APP/PS1*, WSB.*APP/PS1* exhibited lower numbers of parenchymal amyloid plaques but were susceptible to CAA(*9*). Cortical and hippocampal neuronal counts revealed moderate cell loss in both regions in female WSB.*APP/PS1* in comparison with non-carrier littermate controls(*9*).

Here, we describe an extensive characterization of WSB.*APP/PS1* mice, focusing particularly on cerebrovascular phenotyping. Collectively, our data show WSB.*APP/PS1* mice were susceptible to vascular deficits similar to those observed in human ADRD, including CAA and changes to cerebral blood flow and metabolism. The extent of CAA was modified with the addition of humanized *APOE* risk factors. We propose these strains as important preclinical models to uncover mechanisms and test novel therapeutics to treat vascular deficits in ADRD.

## MATERIALS AND METHODS

### Mouse strains and husbandry

All experiments were approved by the Institutional Animal Care and Use Committees (IACUC) at The Jackson Laboratory (JAX: IACUC #12005), Indiana University (IU: IACUC #21150), Allen Institute for Brain Science (IACUC #1908), or University of Oregon (UO: IACUC #AUP-20-25). WSB.*APP/PS1* mice (JR#33567283) were created as described previously(*9*), with all experiments utilizing the *APP/PS1* allele in the hemizygous state. The allelic series of humanized *APOE* was created by backcrossing either *APOE^4^*, *APOE^3^*, or *APOE^2^*(created by MODEL-AD)(*12*), to WSB for at least 6 generations. For these experiments, homozygous humanized *APOE* alleles were utilized. B6 mice were obtained from JAX (JR#000664) and were bred and maintained in the Howell lab colony.

All cohorts of mice were bred and aged in the Howell lab mouse facilities at JAX, maintained in a 12/12-hour light/dark cycle and room temperatures were maintained at 18-24°C (65-75°F) with 40-60% humidity. All mice were housed in positive, individually ventilated cages. Standard autoclaved 6% fat diet, (Purina Lab Diet 5K52) was available to the mice *ad lib*, as was water with acidity regulated from pH 2.5-3.0. Cohorts of mice were either assessed at JAX or shipped to IU for PET/CT imaging, to Allen Insitute for Brain Science for 3D vessel labeling, or to UO for assessment of cerebral artery reactivity/stiffness.

### Immunofluorescence, imaging, and analysis

#### Tissue collection

In accordance with JAX IACUC approval number 12005, mice were anesthetized with 1g/kg ketamine and 200mg/kg xylazine and were transcardially perfused with PBS. Brains were removed from the skull and hemisected. Hemibrains were placed in 4% PFA overnight at 4°C, 15% sucrose overnight at 4°C, and 30% sucrose overnight at 4°C. Hemibrains were frozen and stored at -80°C. Coronal 25µm sections were collected and stored floating in 31.25% glycerol, (Sigma-Aldrich, G5516-1L) 31.25% ethylene glycol (Sigma-Aldrich, 102466) dissolved in PBS at 4°C.

#### Immunofluorescence

Brains sections were washed in PBS, permeabilized with 1% TritonX (Sigma-Aldrich, T8787) dissolved in PBS for 3 10-minute washes. Brains assayed for amyloid plaques were incubated in 40µg/mL X34 (Sigma-Aldrich, SML1954) dissolved in 40% ethanol for 10 minutes, followed by washes in water (3 minutes) and .02M NaOH (5 minutes). Brains were blocked in 5% normal donkey serum (Sigma-Aldrich, D9663) in 1% TritonX PBS for one hour at room temperature. Sections were incubated with primary antibody diluted in blocking solution overnight at 4°C. Primary antibodies used included rabbit anti-IBA1 (Wako, 019–19741, 1.67µg/mL), rat anti-CLEC7A (Invitrogen, mabg-mdect-2, 3.33µg/mL), chicken anti-myelin basic protein (Invitrogen, PA1-10008, 3µg/mL), goat anti-IBA1 (Abcam, ab5076, 5µg/mL), and rat anti-CD68 (Bio-rad, MCA1957, 5µg/mL). Sections were washed in 2% TritonX PBS and incubated in secondary antibodies diluted in 2% TritonX PBS for 2 hours at room temperature. Secondary antibodies included: donkey anti-rabbit 568 (Invitrogen, A10042, 2µg/mL), donkey anti-rat 647 (Abcam, ab150155, 2µg/mL), donkey anti-chicken 647 (Invitrogen, A78952, 2µg/mL), and donkey anti-goat 488 (Invitrogen, A11055, 2µg/mL). Sections were washed in PBS. Sections not assayed with X34 were incubated with 4’,6-diamidino-2-phenylindole (DAPI; Invitrogen, D21490, 1µg/mL) for 5 minutes at room temperature. Sections were mounted onto microscope slides with fluorescent mounting medium (Polysciences, 18606-20). Entire brain sections were imaged at 20x magnification using a Leica DMi8 microscope. All microscope settings were kept identical for each experiment. Images were stitched together in FIJI. Representative ROIs were maximum projections from 25 1µm confocal images taken with an inverted Leica confocal SP8 microscope.

#### Image analysis, CAA grading, and parenchymal plaque quantification

CAA grading was adapted from grading of human sections utilized by the Mayo clinic(*13*), where a brain section received a score of 0 if there was no CAA at all, and 0.5 if there was only leptomeningeal CAA. A section received a score of 1 if there was leptomeningeal CAA and scattered CAA in cortical vessels. A section received a score of 2 if there were leptomeningeal CAA and strong banding of CAA in cortical vessels, and a score of 3 if strong vessel banding was widespread throughout the cortex. For image quantifications, analyses were performed on three coronal brain sections per brain; at approximately bregma -1.955mm, -2.46mm, and -2.88mm. Scores from each of three brain sections were averaged to give the final CAA score for each brain.

Semi-automated plaque quantifications were performed on the same sections as those graded for CAA, using custom FIJI scripts. Brain cortexes were outlined manually using the polygon tool. The cortex ROI was shrunken by 50µm to avoid confounds at the tissue boarder. X34-stained images were subjected to a gaussian blur (sigma = 15) and were thresholded using the triangle method(*14*). Holes were filled and particles 10-50000µm^2^ (an outline slightly larger than individual X34+ plaque areas) were added as ROIs. These ROIs were combined and overlayed onto to the original X34-stained image. Regions within the combined ROI were thresholded using the MinError method(*15*) and particles 100-50000µm^2^ (X34+ plaques) were analyzed for area (plaque size) and were added as ROIs. X34+ cerebral amyloid angiopathy ROIs were manually removed. Number of particles (plaque number) was divided by the cortex ROI area to give X34+ parenchymal plaques/mm^2^.

Plaque ROIs were expanded to give a 30µm ROI from the edge of the plaque. Overlapping expanded ROIs were combined into one ROI. Expanded ROIs were overlaid onto corresponding IBA1- and CLEC7A-assayed images. Regions within the expanded ROIs were thresholded using the triangle method(*14*), and all particles were analyzed for area. Total IBA1+ and CLEC7A+ area per brain section was divided by total plaque area (plaque number*plaque size) in the corresponding brain section to quantify area per X34+ parenchymal plaque area.

### RNA-Sequencing, data processing, and analysis

Left brain hemispheres were collected from male and female WSB.*APP/PS1* mice and wild-type (WT) littermate controls at 4, 8, and 14 months as described for immunofluorescene. Tissues were snap-frozen at the time of harvest and processed by the Genome Technologies Core at JAX for RNA extraction and library preparation as described previously(*16*). Libraries were sequenced in paired-end mode with a target of 40 million read pairs per sample. To ensure even sequencing depth and mitigate lane effects, samples were distributed across multiple flow cell lanes. FASTQ files were later concatenated per sample to produce a single paired-end file for downstream processing.

Raw sequencing reads were initially assessed for quality using FastQC (version 0.11.3, Babraham) and followed by quality-trimming and filtering using Trimmomatic tool (version 0.33) (*17*). High-quality reads were then aligned to a custom *Mus musculus domesticus* (WSB/EiJ strain) reference genome using ‘STAR’ aligner (version 2.5.3)(*18*). The custom genome for the WSB strain was created by integrating REL-1505 variants into the standard mm10 mouse genome(*19*). To enable detection of the human *APP/PS1* transgenes, we also appended sequences of *APP* and *PSEN1* to the reference genome and annotation files. Gene expression was quantified using two complementary approaches to support multiple downstream analytical methods: transcripts per million (TPM) using RSEM (version 1.2.31)(*20*) and raw read counts using HTSeq count (version 0.8.0)(*21*). Expression of *APP* and *PSEN1* transgenes was confirmed by inspecting RNA abundance in the engineered mouse model.

To reduce noise and improve statistical power, lowly expressed genes were filtered out by requiring a minimum total count across the samples. Genes with zero counts across all samples or have less than 10 reads in 3 or more samples were filtered out. For sex-adjusted analyses, genes located on the X and Y chromosomes were removed prior to PCA and DE modeling to avoid sex-chromosome-driven variation. Normalization and differential expression analysis were carried out using DESeq2(*22*), accounting for sex, age and genotype effects. Variance-stabilizing transformation (VST) was applied to normalized counts for downstream exploratory analyses and visualization. Principal Component Analysis (PCA) was used to assess sample-level clustering and detect potential outliers. All raw and processed is being made available through the AD Knowledge Portal.

#### Functional enrichment analysis

Transcriptional profiling aimed to characterize genotype-associated transcriptional alterations across different ages in the WSB background. Analyses were stratified into three comparisons: genotype effect at 4, 8, and 14 months, based on sex-adjusted and sex-specific models. DESeq2 was used to identify gene expression differences between experimental groups, and the results were exported as log2 fold changes and associated statistics.

For each comparison, Gene Set Enrichment Analysis (GSEA)(*23*) was performed on pre-ranked gene lists (ordered by log2 fold change) using the gseGO(), gseKEGG(), and gsePathway() functions from the clusterProfiler and ReactomePA packages. Gene symbols were mapped to Entrez IDs via org.Mm.eg.db. Enrichments were computed separately for Biological Process (BP) Gene Ontology terms, KEGG pathways, and Reactome pathways. For exploratory analysis, significantly enriched pathways were defined based on normalized enrichment scores (NES) and adjusted p-values (< 0.2).

Selected biologically relevant pathways were visualized using customized gene-pathway network plots. For each pathway, core enriched genes identified by GSEA were mapped to their associated terms, allowing visualization of gene-level contributions within each module and experimental condition.

#### Mouse-human transcriptomic mapping

To evaluate the relevance of transcriptional changes in WSB mice to human AD, we utilized gene co-expression modules derived from the Accelerating Medicines Partnership for Alzheimer’s Disease (AMP-AD) consortium. A total of 30 human brain co-expression modules were obtained from the Synapse data repository (SynapseID: syn11932957), originally constructed by Wan and colleagues(*24*) through a meta-analysis of differential gene expression across seven brain regions from three independent late-onset AD (LOAD) cohorts (*25–27*). These modules were further clustered into five functional consensus groups representing major transcriptomic alterations in human late-onset AD.

We assessed the degree of concordance between genotype-driven expression changes in WSB mice and AMP-AD modules using a fold-change correlation approach(*28*). Specifically, we examined both sex-specific and sex-adjusted genotype effects at 4, 8, and 14 months of age, yielding multiple biologically distinct comparison groups. For each of these comparisons, Pearson correlation coefficients were calculated between the log2 fold-change values for mouse genes (relative to WT controls) and the corresponding log2 fold-change values observed in human AD cases versus controls. Human transcript-level fold-change data were obtained from the AD Knowledge Portal (SynapseID: syn14237651). Each AMP-AD module was assessed for concordance with each age- and sex-stratified genotype effect using the correlation coefficient and its associated p-value.

Results were visualized as module–condition correlation matrices plotted as bubble charts. Only significant correlations (*p <* 0.05) were shown with blue representing positive and red representing negative correlations. Circle size and color intensity reflected the magnitude of the Pearson correlation coefficient.

#### Assessment of mouse-human shared gene signatures

To further investigate the biological relevance of transcriptional overlaps between WSB mouse models and human AD, we focused on the subset of genes that contributed to significant correlations with AMP-AD modules (“intersecting gene sets”). These gene sets were extracted based on their similar differential expression pattern in both species.

Overrepresentation analysis (ORA) was performed using the enrichGO() function from the clusterProfiler R package to identify enriched GO-BP terms within these intersecting gene sets. The top enriched GO terms for each comparison–module pair were visualized using bar plots, ranked by adjusted p-value. These plots highlight biological processes most consistently altered across species. In addition, we performed GSEA using gseGO() for these intersecting genes ranked by log2 fold-change in mouse models. Results were visualized using split bar plots to display the top up- and down-regulated pathways (based on NES), with bar lengths proportional to NES and colored by gene set size. To capture the gene-level contributions to specific pathways, we constructed bipartite gene–pathway networks for top enriched terms (from GSEA) using the igraph package. Separate networks were generated for upregulated and downregulated pathways. Term and gene nodes were color-coded by directionality (e.g., red/green for up, blue/orange for down), and networks were visualized using force-directed layouts to aid interpretability.

### PET/CT Imaging and Analysis

PET/CT imaging protocols were performed in accordinace with IU IACUC approval number 21150. Evaluation of metabolism and perfusion was as described previously(*16*).

#### Radiopharmaceuticals

Regional brain glycolytic metabolism was monitored using 2-[^18^F]-fluoro-2-deoxy-D-glucose (^18^F-FDG), which was synthesized, purified, and prepared according to established methods, where unit doses (185 to 370 MBq) were purchased from PETNet Indiana (PETNET Solutions Inc). Brain perfusion was measured using Copper(II)-[^64^Cu]-pyruvaldehyde-bis(N4-methylthiosemicarbazone) (^64^Cu-PTSM), which was synthesized, purified, and unit doses (i.e. 370 to 740 MBq) dispensed by the PET Radiochemistry Core Facility at Washington University according to methods described previously(*29, 30*).

#### Positron Emission Tomography (PET) and Computed Tomography (CT) Imaging

To evaluate changes in cerebral glycolysis and cerebral perfusion mice were placed in a restrainer and consciously injected into the peritoneal cavity (^18^F-FDG) or tail vein (^64^Cu-PTSM) with 0.24-3.04 MBq of purified, sterile radiotracer, where the final volume did not exceed 10% of the animal’s body weight. Each animal was returned to its warmed home cage and allowed 30 minutes (^18^F-FDG) or 5 minutes (^64^Cu-PTSM) to allow for uptake and cellular trapping(*30, 31*). Animals were fasted overnight only for imaging with ^18^F-FDG. Post-uptake, mice were induced with 5% isoflurane gas (95% medical oxygen), placed on the imaging bed, and anesthesia maintained at 1-3% isoflurane (97-99% medical oxygen) during acquisition, per our previous work(*16*). To provide both anatomical and function images, PET/CT acquisitions were performed with a Molecubes β-X-CUBE system (Molecubes NV, Gent Belgium), where calibrated list-mode PET images were acquired in list mode for 10 (^18^F-FDG) or 20 (^64^Cu-PTSM) minutes, and reconstructed into a single-static image using ordered subset expectation maximization (OSEM) with 30 iterations and 3 subsets(*32*). To provide anatomical reference and attenuation maps necessary to obtain fully corrected quantitative PET images, helical CT images were acquired with tube voltage of 50 kV, 100 mA, 100 μm slice thickness, 75 ms exposure, and 100 μm resolution. For β-CUBE studies, images were corrected for radionuclide decay, tissue attenuation, detector dead-time loss, and photon scatter previously described(*32*).

#### PET/CT Image Processing and Analysis

All PET/CT images were co-registered using a ridged-body mutual information-based normalized entropy algorithm(*33*) with 9 degrees of freedom and mapped to stereotactic mouse brain coordinates(*34*) using MIM Encore Software 7.3.2 (Beachwood OH). Study imaging and demographics data was managed using RedCap database hosted at Indiana University. Post-registration, 56 bilateral atlas regions were extracted, and left/right averaged to yield 27 unique volumes of interest that map to key cognitive, sensory, and motor centers. To permit dose, scanner and brain uptake normalization, Standardized Uptake Value Ratios (SUVR) relative to the cerebellum were computed for PET for each subject, genotype, age, and condition as follows:

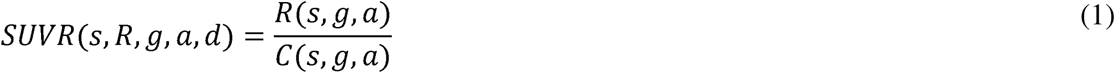

where, *s, g, a, R*, and *C* are the subject, genotype, age, region/volume of interest, cerebellum region/volume of interest. The SUVR values were then converted to z-score as follows:

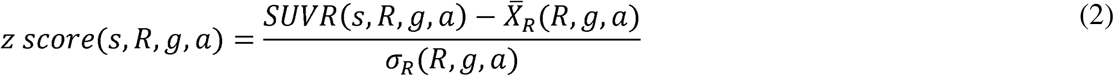

where, *s, g, a, R*, *X̄_R_* and *σ_R_* are the subject, genotype, age, mean of the reference population in SUVR, standard deviation of the reference population, based on the specified analytical strategies (effects of diet, aging, humanized genes, and AD-risk alleles). Data are then projected onto Cartesian space per our previous work(*4*), where the x-axis represents the z-score change in perfusion, derived from the ^64^Cu-PTSM data, and the y-axis is the z-score change in glycolytic metabolism(*35–37*) via ^18^F-FDG.

To determine the radiotracer kinetics, arterial input function (*C_a_*) were derived from whole blood measurements of ^64^Cu-PTSM and were digitized and intensity normalized. At steady-state, ^64^Cu-PTSM is trapped in the tissues directly proportional to tissue perfusion(*30*), where estimates of the tissue conductance is governed by the permeability-surface area product (*PS*) in a one tissue compartment model. Model estimates were determined using the autoradiographic approach(*31, 38, 39*), by solving the following partial differential equation:

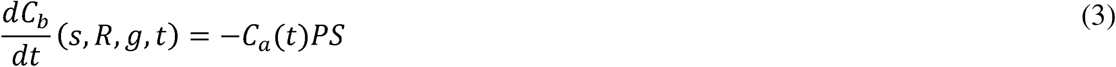

where, *s, g, R, t, C_b_*, and *C_a_* are the subject, genotype, region/volume of interest, time, concentration in the brain, and arterial input function. Estimates of PS were determined iteratively, using an average dose of 4.63 MBq and 2.8ml average blood volume. Parameters were optimized via Levenberg-Marquardt algorithm using a sum-of-squares error of the absolute relative difference between the measured tissue tracer levels and estimated final value based on:

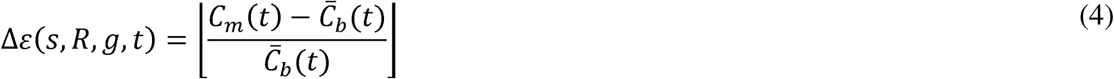

where, Δ*ε, s, g, R, t*, *C̄_b_* and *C_m_* are the absolute error difference, subject, genotype, region/volume of interest, time, average brain concentration, and modelled concentration, respectively. In all cases, trapezoidal quadrature was used to numerically integrated *C_a_* with time using Eqn. 3, where a tolerance of 0.01 was used as the stopping criteria. The error estimate of this model was 0.95±0.033% with an averaged R^2^ of 0.9984 across all subjects, genotypes, and regions.

### 3D Vessel Labeling, Imaging, and Quantification

3D vessel labeling and imaging were performed in accordance with Allen Insitute for Brain Science IACUC approval number 1908. A separate cohort of 8-month old male and female WSB.*APP/PSI* mice were shipped to the Allen Institute for Brain Science and allowed to acclimate for 72 hours before handling. Twenty-four hours prior to euthanasia, all animals received an intraperitoneal injection of 3.3mg/kg methoxy-X04 in 3.3% DMSO, 6.7% Kolliphor-EL (Millipore Sigma) in PBS. Blood vessel lumen staining was performed using the protocol described previously(*40*). At harvest, animals were perfused with 10mL of 0.9% NaCl, followed by 50mL of 4% PFA. After PFA perfusion, the mice were tilted ∼30°, head down, with their head placed in ice water. The carcass was then perfused with 20ml of heated 2% porcine skin gelatin (Sigma 1890) and 0.1% FITC-albumin (Sigma A9771), then the heart was clamped to stop fluid circulation and the head submerged in ice water for 30 minutes to allow the gelatin to solidify. The brain was carefully removed from the skull and placed into post-fix solution overnight, then sectioned and imaged as described previously(*41*). To provide quantification of vascular tree volume, exchange surface area, and permeability, stacks of 140 32bit Red/Green/Blue/Alpha (RGBA) images were used to reconstruct a 3D volume using cubic spline interpolation, where the slice offset was based on the image section thickness (100μm). The 16-bit transformed green channel contained the information required to detect, visualize, and extract 3D vessels. Post-reconstruction, image volumes were registered to the Allen Common Coordinate Framework(*42*) permitting alignment to a common reference space.

Due to the variance between voxel intensity values and subjects, a method to automatically normalize and detect the precise range of vessel values was developed and utilized the entropy of the volume for different ranges of voxel intensities that spanned from 0 to 65535. Data plotted as entropy vs. intensity range were used to determine the point where the entropy curve starts to decrease (i.e. Δ>0.01), which was selected as the minimum of the threshold range (**Fig. S1A**).

This threshold is used to obtain a coarse initial segmentation of the vessels’ volume. Then, to establish the 3D connection between vessels, we used a 26-neighbors connected components process to reduced spurious voxels in the image volume that were not part of the vascular tree.

Once the vessels were extracted, the vascular tree was skeletonized by applying the medial axis transform(*43*), thus providing a midline throughout the entire tree that could be used for additional processing. This process facilitated the counting of the number of vascular branches, since the process was reduced to finding the locations where the vessels split, and branches are defined as a segment, or connected voxels, between two splitting locations (**Fig. S1B**). To perform this task, the algorithm started at any voxels that belong to the skeletonized vessels’ tree, then traverses the complete tree until all voxels are explored via a divide and conquer approach. Traversing the vessels tree was defined as following the path of connected voxels and enumerating each voxel where the path divides (**Fig. S1B**). To prevent the algorithm from cycling infinitely, our method keeps track of which voxels were visited during this process.

To evaluate the vessel diameters for each branch, the algorithm iterated across all detected branches, and was fitted with a polynomial along the branch voxels to permit interpolation of elements. A normal vector between consecutive voxels of the fitted curve was used to obtain an orthogonal plane placed between the voxels being evaluated. Using the orthogonal plane and the normal vector, a 2D slice projection of the vascular tree was generated. In combination with the completed vascular tree, a ray casting process is used to calculate the vascular exchange surface (perimeter of the slice) and cross-sectional areas assuming a prolate ellipsoid, where origin of the ray was centered along the midline, and the ray in each direction intersects the vessel contour 180 out of phase. This process is repeated iteratively by rotating the angle between rays over the interval of 0° to 180° with 1° steps (**Fig. S1C**). The algorithm returns a set of measurements for each voxel in the skeletonized vessel tree.

### Cerebral artery reactivity

#### Tissue collection

In accordance with University of Oregon IACUC approval number AUP-20-25, mice were euthanized via isoflurane (5%) overdose, followed by exsanguination by cardiac puncture and a bilateral thoracotomy. Vasodilation and vasoconstriction responses were assessed *ex vivo* in isolated, pressurized posterior cerebral arteries, as described in detail(*44, 45*). Briefly, arteries were excised from the brain and placed in pressure myograph chambers (DMT Inc., Hinnerup Denmark) with physiological salt solution containing 145 mM NaCl, 4.7 mM KCl, 2 mM CaCl_2_, 1.17 mM MgSO_4_, 1.2 mM NaH_2_PO_4_, 5.0 mM glucose, 2.0 mM pyruvate, 0.02 mM EDTA, 3.0 mM MOPS buffer, 10 g/L BSA, 7.4 pH at 37°C. The arteries were then cannulated onto glass micropipettes and secured with nylon (11–0) sutures. Once cannulated, arteries were warmed to 37°C, pressurized to 50 mmHg, and equilibrated for approximately 1 hour. All arteries were submaximally preconstricted with phenylephrine (1-6 μM to obtain ∼15-40% preconstriction).

#### Vasoreactivity measurements

To measure vascular reactivity, changes in lumen diameter were measured in response to increasing concentrations of endothelium-dependent vasodilator acetylcholine (1×10^−9^ to 1×10^−4^ M), endothelium-independent vasodilator sodium nitroprusside (1×10^−10^ to 1×10^−4^ M), or vasoconstrictor endothelin-1 (1×10^−11^ to 1×10^−7^ M). Acetylcholine dose responses were repeated after incubation with L-N^G^-nitro arginine methyl ester (L-NAME, 0.1 mM for 30 minutes) to examine the contribution of nitric oxide synthase to endothelium-dependent vasodilation.

#### Cerebral artery stiffness

Passive arterial stiffness was measured *ex vivo* in the posterior cerebral artery by the resulting lumen diameter and medial wall thickness following increasing intraluminal pressure(*44*). Arteries were isolated and cannulated between glass pipette tips as described in “Tissue Collection”. Passive stiffness was measured in arteries following 60-minute incubation in a calcium-free solution to eliminate the effects of myogenic tone. Measurements for each artery were recorded from 5 to 100 cmH_2_O (3.7–73.5 mmHg) in 5 cmH_2_O increments. Stress-strain curves were created for each artery to calculate β-stiffness. Stress was calculated as: σ = PD/2WT, where P is pressure in dyne cm^-2^, D is lumen diameter and WT is wall thickness. Strain was calculated as: ε = (D - D_i_)/D_i_, where D_i_ is the initial lumen diameter. Data for each artery were fit to the curve: σ = σ_i_e^βε^, where σ_i_ is the initial stress (at 5 cmH_2_O), and β is the slope of tangential elastic modulus versus stress. A higher β-parameter represents a stiffer artery.

### Statistical analysis

Data were analyzed using GraphPad Prism9 software. Power calculations were performed before experiments were conducted to determine appropriate sample size. Data from experiments designed to test differences between two groups were subjected to an *F* test to compare variance and a Shapiro–Wilk test to test normality to ensure appropriate statistical tests were utilized. Normally distributed data with similar variance were analyzed using unpaired two-tailed *t* tests. Non-normally distributed data were analyzed using Mann-Whitney U tests. Data from experiments designed to test differences among more than two groups across one condition were subjected to a Brown-Forsythe test to compare variance and a Shapiro–Wilk test to test normality to ensure an appropriate statistical test was utilized. Normally distributed data with similar variance were analyzed using a one-way ANOVA with Tukey’s post-hoc test. Non-normally distributed data were analyzed using a Kruskal–Wallis test with Dunn’s post hoc test. Normally distributed data with unequal variances were analyzed using a Brown-Forsythe ANOVA with Dunnett’s T3 multiple comparisons test. Data from experiments designed to detect differences among multiple groups and across two conditions were analyzed using a two-way ANOVA with Holm-Sidak’s post-hoc test. Data from experiments designed to test differences between two groups across multiple conditions were analyzed with a three-way ANOVA. For these statistical tests, every possible comparison was made, and multiplicity adjusted *P* values are reported. In all cases, data met the assumptions of the statistical test used and *p* values < 0.05 were considered statistically significant. Male and female mice were treated as individual biological groups and were analyzed separately in all cases.

## RESULTS

We have previously shown WSB.*APP/PS1* develop cognitive deficits in working memory (as determined in the spontaneous alternation assay), limited but significant neuronal cell loss, and increased instances of CAA compared to B6.*APP/PS1* mice(*46*), suggesting the WSB genetic context may be more conducive to study the vascular contributions to ADRDs. Therefore, we first set out to better characterize the WSB.*APP/PS1* strain, focusing particularly on the anatomy and function of the cerebral vasculature.

### WSB.APP/PS1 developed human-relevant CAA

To characterize CAA in WSB.*APP/PS1* mice, 4-, 8-, and 14-month-old (M) female and male WSB.*APP/PS1* mice were evaluated for CAA and parenchymal amyloid plaque deposition (**Fig. 1**). At 4M, few X34+ parenchymal plaques were observed, and no vascular X34+ CAA was observed in either sex or genotype. Both male and female WSB.*APP/PS1* mice developed increasingly severe CAA (**Fig. 1C**) and parenchymal plaque deposition (**Fig. 1D**) at 8 and 14M. Plaque size remained consistent with age for each sex (**Fig. 1E**).

**Figure 1:**
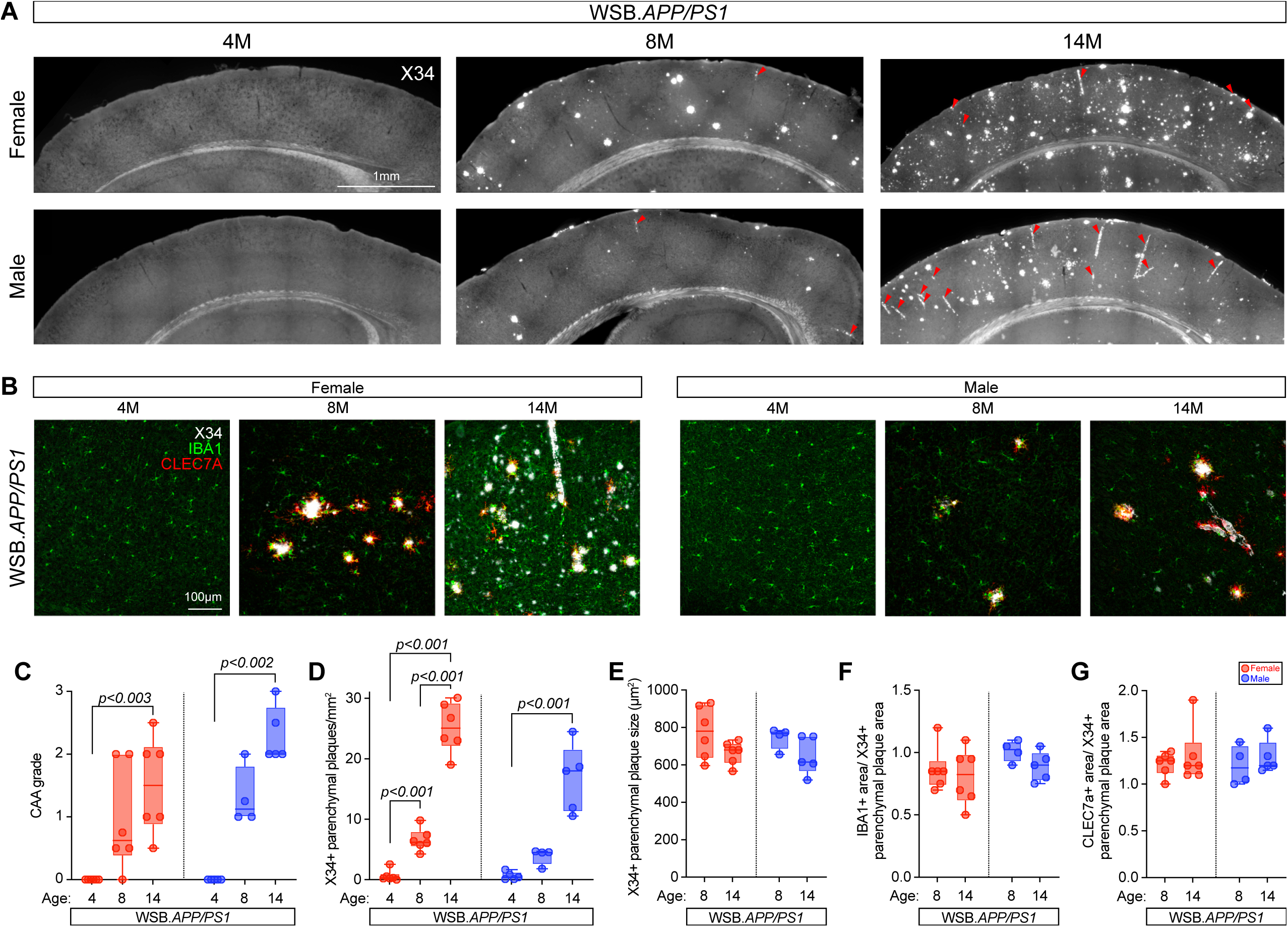
CAA and parenchymal plaque characterization in WSB.APP/PS1 mice. **(A)** Female and male 4M, 8M, and 14M WSB.*APP/PS1* brain hemisections stained with X34 (white) and immunoassayed for IBA1(green) and CLEC7A (red). Arrowheads indicate CAA. Magnification: 20X; scale bar: 1mm. **(B)** High-resolution insets of representative parenchymal plaques and CAA. Magnification: 40X; scale bar: 100µm. **(C)** Quantification of CAA scores. CAA scores significantly increased with age in both sexes (*p <* 0.001, Kruskal-Wallis tests, Dunn’s multiple comparisons tests). Magnification, 40X; scale bar, 100µm. **(D)** Cortical parenchymal X34+ plaque quantifications normalized to cortical area. Both male and female WSB.*APP/PS1* had increased parenchymal plaque deposition with age (female: *p <* 0.001, Brown-Forsythe ANOVA test, Dunnett’s T3 multiple comparisons test; male: *p <* 0.001, Kruskal-Wallis test, Dunn’s multiple comparisons test). **(E)** Average plaque size quantifications. No differences were observed in either sex across ages (female: *p =* 0.107; male: *p =* 0.146; two-tailed t-tests). **(F-G)** Average IBA1+ area **(F)** and CLEC7A+ area **(G)** normalized to average plaque size. No differences were observed across age for IBA1+ (*p =* 0.629 and *p =* 0.129 for female and male respectively, two-tailed t-tests) or CLEC7A+ area (female: *p =* 0.831, Mann-Whitney test; male: *p =* 0.514, two-tailed t-test). For 4M, 8M, and 14M, respectively; female: *n =* 6, 6, 6; male: *n =* 5, 4, 5. Female data indicated in red, male data in blue. (C-G).

Previous studies have demonstrated that microglial activity modifies CAA and parenchymal plaque development in mice(*47*). Moreover, we have previously shown WSB mice have fewer microglia compared B6(*9*), and WSB microglia were shown to respond differently to amyloid compared to other strains(*10*). Therefore, plaque-associated microglial (IBA1+) and CLEC7A+ (a known disease-associated microglial protein(*48–50*)) area was measured within a 30µm radius from each X34+ parenchymal plaque. As anticipated, there was robust IBA1+ and CLEC7A+ expression in proximity to X34+ parenchymal plaques, and this expression remained consistent across ages and sexes (**Fig. 1F, G**).

### Neuronal, vascular, and immune related transcriptional changes emerged progressively in WSB.APP/PS1 mice

To identify potential mechanisms important in driving AD-relevant phenotypes, we employed transcriptomic profiling of brain tissues from male and female WSB and WSB.*APP/PS1* mice at 4, 8, and 14M. These analyses revealed distinct temporal patterns in biological pathways as early as 4M (**Fig. 2A**), before overt amyloid plaque deposition (**Fig. 1**). At this early stage, enriched pathways were dominated by neuron projection organization and regulation of synaptic plasticity, with upregulation of genes such as *Xlr3b*, *Kdr* and *Rims1*. Upregulation of *Rims1*, a presynaptic scaffolding protein, supports neurotransmitter release and short-term synaptic plasticity(*51*), suggesting early compensatory remodeling of neuronal circuits. *Kdr* encodes VEGFR2, a major endothelial growth factor receptor essential for neurovascular coupling and angiogenesis(*52, 53*), and also plays a neuroprotective role. These neuronal changes are accompanied by transcriptional signatures of glial cell activation, indicating that neuroinflammatory processes are initiated prior to detectable amyloid pathology in WSB.*APP/PS1*. Notably, genes related to response to starvation and the endoplasmic reticulum (ER) unfolded protein response (UPR) were downregulated, including *Xbp1* and *Hspa5*. *Xbp1* is a master transcription factor in the ER stress response and regulated metabolic adaptation to nutrient limitation(*54*), while *Hspa5* encodes a central ER chaperone maintaining protein folding homeostasis, which is involved in memory consolidation(*55*). Downregulation of the UPR signaling pathways has been linked to *PS1* mutations(*56*) and potentially impair the ability of neurons and vascular cells to buffer misfolded proteins and contribute to vulnerability in preclinical AD stages. Chronic low-level ER stress has been implicated in promoting vascular endothelial dysfunction via oxidative stress and inflammatory signaling(*57*). In this context, early molecular changes in nutrient-sensing and proteostatic pathways could represent an upstream trigger for the vascular and inflammatory signatures observed at later ages.

**Figure 2:**
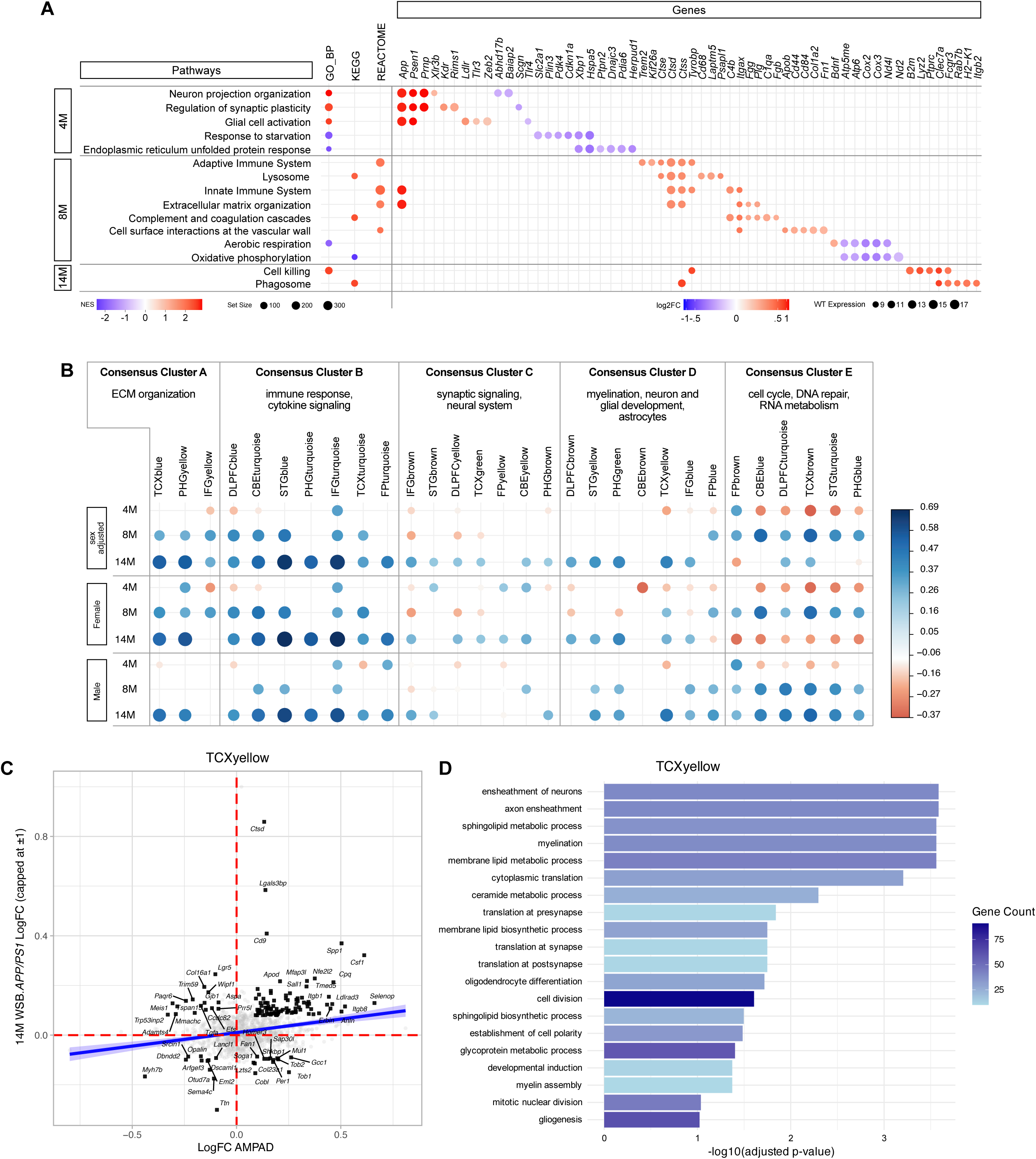
Transcriptional profiling in WSB and WSB.APP/PS1 mice. Gene set enrichment analysis in WSB.*APP/PS1* mice (compared to WSB) at 4, 8 and 14M. (**A**) Age-dependent pathways identified by GO, KEGG and Reactome. Genes relating to each pathway are shown to the right. For pathways, positive NES scores are shown in red, negative NES scores are shown in blue. For genes, increased log2 fold change is shown in red, decreased log2 fold change is shown in blue. (**B**) Alignment of differentially expressed genes in mouse with differentially expressed genes in human AMP-AD modules. A positive correlation is shown in blue, anti-correlation is shown in red. First three rows are combined for sex, second set of three rows are female only, third set of rows are male only. (**C**) One module, TCX yellow in Consensus Cluster D showed age-dependent correlations (strongest correlations at 14M), and included genes involved in microglia function and myelination. (**D**) Enrichment of genes positively correlated in human AMP-AD and WSB.*APP/PS1* mice identified myelination-related processes, predicting white matter changes.

By 8M, the transcriptional profile shifted toward a strong vascular–immune axis (**Fig. 2A**). Significant enrichment was observed in pathways increasingly recognized as central to CAA and small vessel disease in AD, such as extracellular matrix (ECM) organization, complement and coagulation cascades, and cell surface interactions at the vascular wall. Among the upregulated genes, *Itgax* marks a subset of activated microglia enriched in neurodegenerative contexts(*58*), while *Cd44* mediates vascular inflammation and blood-brain barrier dysfunction(*59*). These signatures, coupled with downregulation of aerobic respiration and oxidative phosphorylation, point to a convergence of vascular injury, metabolic compromise, and immune activation, consistent with core features of CAA-associated pathology in AD(*60*).

At 14M (**Fig. 2A**) transcriptional changes were dominated by microglial activation and phagocytic clearance. Enriched immune pathways included cell killing (*Tyrobp*, *B2m*, *Clec7a*, *Lyz2*, *Ptprc*) and phagosome (*Ctss*, *Clec7a*, *Fcgr3*, *Itgb2*, *Rab7b*). *Tyrobp* is a core hub in TREM2-dependent microglial activation networks in AD(*61*), while *Clec7a* marks disease-associated microglia clustered around amyloid deposits(*62*). The upregulation of these pathways and genes is consistent with sustained neuroinflammation and heightened immune mediated clearance activity in advanced amyloid disease.

Overall, the temporal cascade in WSB.*APP/PS1*, beginning with early neuronal plasticity changes at 4M, progressing through vascular and ECM remodeling at 8M, and culminating in immune–metabolic activation under high amyloid load at 14M, aligns with recent human and preclinical evidence supporting metabolic and vascular dysfunction as an early driver of downstream neurodegeneration in AD(*63*).

### Correlation of WSB.APP/PS1 transcriptional profiles with human AMP-AD modules across age and sex

To evaluate the AD relevance of age- and sex-specific transcriptional changes in WSB.*APP/PS1*, we computed correlations between mouse differential expression profiles and 30 human AMP-AD co-expression modules (**Fig. 2B**)(*24*). In terms of human age equivalence, 4, 8, and 14M mice roughly correspond to early adulthood (∼20–30 years), middle age (∼35–45 years), and late middle age to early elderly (∼55–65 years) in humans, respectively(*64*). This framework enables interpretation of transcriptomic shifts in WSB.*APP/PS1* mice along a compressed timeline, mapping them onto the stages of AD pathophysiology observed in humans.

At 4M, correlations were generally modest, reflecting the limited overlap between early-stage molecular alterations in WSB.*APP/PS1* mice and the largely late-stage signatures represented in the AMP-AD cohort. Nonetheless, certain neuronal and metabolic modules exhibited weak but significant correlations, suggesting that subtle network remodeling and early metabolic shifts are already underway.

Correlations strengthened markedly for immune–microglial and vascular/ECM-related modules at 8M (**Fig. 2B**), consistent with the emergence of neuroinflammatory activation and cerebrovascular remodeling during mid-stage disease(*65*). These changes mirror the vascular amyloid deposition and immune cell recruitment characteristic of WSB.*APP/PS1* pathology. At the same time, significant but weak correlations start to emerge for neuronal, synaptic and glial modules in both positive and negative directions. This pattern likely reflects the complex cellular remodeling processes occurring at this stage.

By 14M (**Fig. 2B**), correlations with human AMP-AD modules became more pronounced and broadly distributed across multiple functional domains. The highest positive correlations were seen for consensus clusters A and B (modules enriched for ECM organization, immune activation, and cytokine signaling) consistent with the pronounced vascular remodeling and neuroinflammatory signatures that dominate this late stage. Cluster D, spanning myelination, neuronal and glial development, and astrocyte biology, also showed significant positive correlations, indicating glial activation and changes in neuroglial support networks under sustained amyloid burden. Cluster C, representing synaptic signaling and broader neuronal system pathways, displayed more significant correlations in females than in males, with limited overlap between the sexes at the individual module level. This sex-biased divergence suggests that synaptic remodeling programs are differentially engaged in males and females during advanced disease. Finally, Cluster E, which includes cell cycle, DNA repair, and RNA metabolism modules, exhibited striking sex-specificity: correlations were in opposite directions between males and females, pointing to fundamentally different regulation of cell cycle and genome maintenance pathways in the aging mouse brain depending on sex.

Overall, the AMP-AD module mapping revealed distinct temporal trajectories in how WSB.*APP/PS1* transcriptional profiles align with human AD signatures. Vascular, immune, and ECM–related modules showed a progressive strengthening of positive correlations from mid- to late-stage disease, suggesting these processes represent a sustained and converging pathology. Myelination and astrocyte related modules became more prominent at later stages, while synaptic signaling modules displayed a marked sex bias, with stronger and more numerous correlations in females. The most striking sex specificity emerged in modules related to cell cycle, DNA repair, and RNA metabolism, where the direction of correlation reversed between males and females. Collectively, these patterns highlight that the temporal evolution of molecular networks in WSB.*APP/PS1* mice is shaped both by disease stage and by sex, with some processes showing steady progression and others exhibiting divergent, sex-dependent trajectories.

### WSB.APP/PS1 mice exhibited human-relevant white matter changes

White matter damage is often a key indicator of cerebrovascular damage in human patients. Interestingly, the TCXyellow module from the consensus cluster D, representing myelination, neuronal and glial development, and astrocyte biology, showed strong positive correlations between WSB.*APP/PS1* sex-adjusted expression profiles at 14M and human AD transcriptional changes (**Fig. 2B**). The key contributing genes driving the strong mouse–human correlation included *Spp1* and *Csf1* (key mediators of microglia–oligodendrocyte crosstalk and glial activation), *Cpq* and *Apod* (linked to lipid metabolism and myelin turnover), and *Nfe2l2* and *Itgb1* (stress response and ECM signaling) (**Fig. 2C, D**). This pattern suggests the human–mouse overlap reflects an integrated glial response involving astrocytic, microglial, and vascular-interacting pathways.

GO enrichment analysis of the TCXyellow module at 14M revealed strong overrepresentation of biological processes involved in myelin assembly, oligodendrocyte differentiation, and axon ensheathment. Additional enriched terms included lipid and sphingolipid biosynthetic processes, as well as translation at pre-, post-, and general synaptic compartments, suggesting concurrent remodeling of myelin and synaptic machinery. The enrichment of both myelin-related and lipid biosynthesis pathways is consistent with the high metabolic demands of oligodendrocyte maintenance and repair(*66*) that would be predicted to result in myelin changes in WSB.*APP/PS1* mice. To further validate this, sections that included the corpus callosum and cortex from 14M male and female WSB.*APP/PS1* and WSB control mice were stained with antibodies to visualize myelin (myelin basic protein, MPB, **Fig. 3**) and MBP intensity evaluated. Compared to WSB controls, WSB.*APP/PS1* showed a significant reduction in MBP intensity in the corpus callosum of male and female mice, and in the cortex of male mice (**Fig. 3C, D**). The reduction of cortical MBP intensity in male mice aligned with the stronger correlation of the TCXyellow module in male compared to female mice (**Fig. 2D**). Collectively, these data support human-relevant white matter damage in WSB.*APP/PS1* mice.

**Figure 3:**
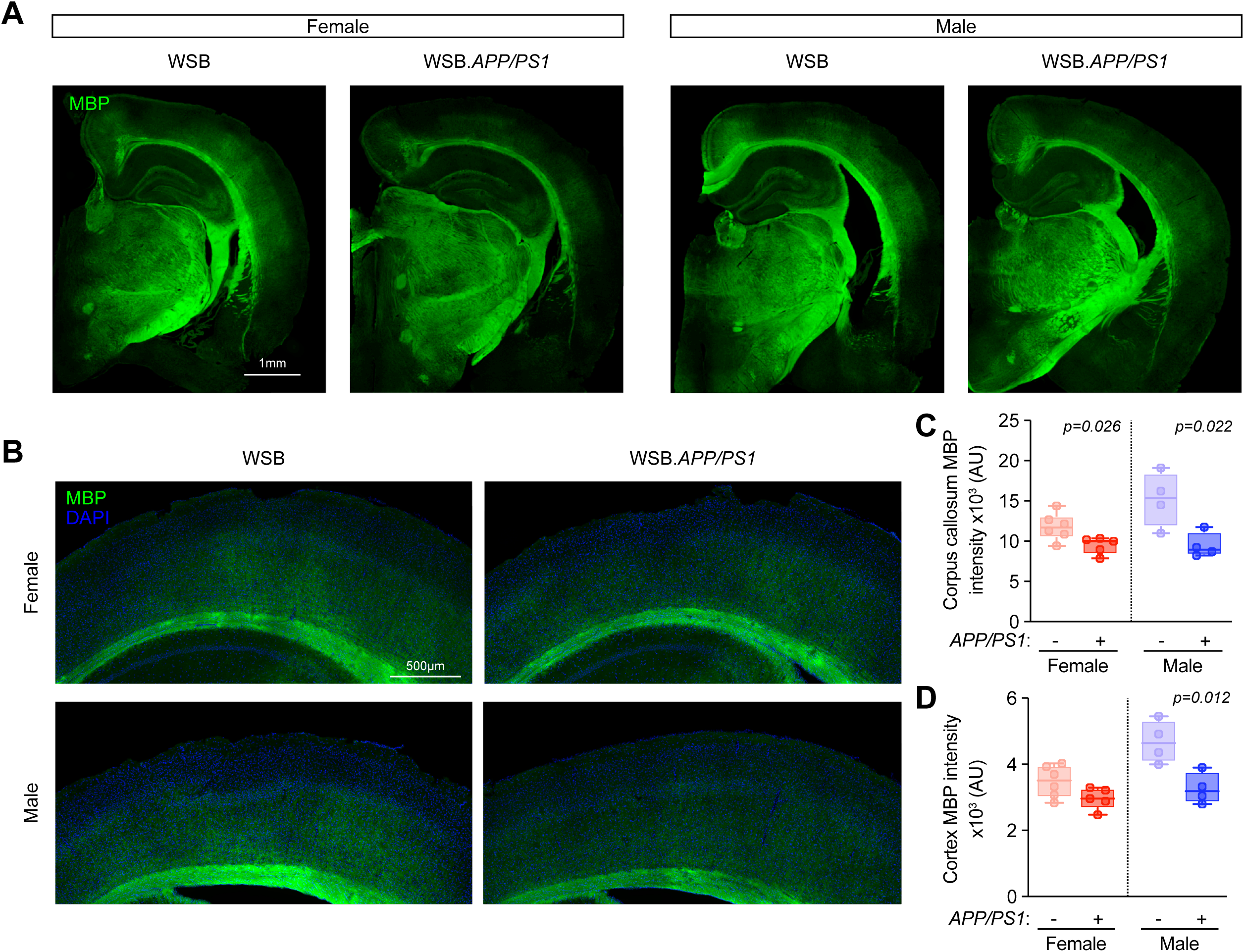
White matter changes in WSB.APP/PS1 mice. (**A**) Representative 14M female and male WSB WT and WSB.*APP/PS1* brain hemisections immunoassayed for myelin basic protein (MBP, green). Magnficiation: 20X; scale bar: 1mm. (**B**) Representative regions of interest counterstained with DAPI (blue) Magnification: 20X; scale bar: 500µm. (**C-D**) Quantification of corpus callosum **(C)** and cortical **(D)** MBP intensity. Male WSB.*APP/PS1* had significantly reduced MBP intensity in both the corpus callosum and the cortex compared to WT controls (*p =* 0.022 and 0.012, respectively; two-tailed *t* tests). Female WSB.*APP/PS1* mice had a significant reduction in MBP intensity in the corpus callosum (*p =* 0.026, two-tailed *t* test) but did not have a statistically significant decline in MBP intensity in the cortex (*p =* 0.072, two-tailed *t* test) compared to WT controls. For WT and *APP/PS1*, respectively: female: *n =* 6, 5; male: *n =* 4, 4.

### WSB.APP/PS1 showed vascular and metabolic uncoupling, increased vascular tree volume, and surface area changes

To understand the relationship between cerebral blood flow (CBF) and metabolism, we performed regional neurovascular uncoupling analysis per our previous work(*4*). As anticipated, WSB.*APP/PS1* relative to WSB showed significant regional metabolic and vascular dysregulation (MVD) (**Fig. 4A, B**), with most brain regions in both sexes showing prodromal (PD, increased perfusion, increased metabolism) or neuro-metabolic and vascular failure (NMVF, decreased perfusion, decreased metabolism) coupled phenotypes. In addition to these changes, WSB.*APP/PS1* mice showed a divergent sexually dimorphic regional pattern. Male WSB.*APP/PS1* mice (relative to male WSB WT) showed Type 1 uncoupling (T1U, increased perfusion, decreased metabolism) in the agranular insular cortex (AI), ectorhinal cortex (ECT), and fornix, while the lateral orbital (LO), ventral orbital (VO), hippocampus (HIP), and prelimbic (PrL) showed Type 2 uncoupling (T2U, decreased perfusion, increased metabolism) (**Fig. 4A, B**). Moreover, the dorsolateral-inferior-ventral entorhinal (DLIVEnt) and visual 1 and 2 (V1V2) cortices showed the PD phenotype, while the parietal association cortex (PtA) is in NMVF. By contrast, female WSB.*APP/PS1* (relative to female WSB WT) mice showed significant T2U in temporal association cortex (TeA), parietal association cortex (PtA), and secondary sensory cortex (S2), while the Perirhinal Cortex (PRH) showed a PD phenotype (**Fig. 4A, B**). To understand if these patterns were related to changes in vascular permeability (P) and surface area (S) changes, we kinetically modeled the ^64^Cu-PTSM images, yeilding regional PS product (ml/g.min), which is a measure of the tissue bulk permeability (**Fig. 4C, D**). Females had generally higher PS compared to males irrespective of genotype. When sexes were analyzed separately, compared to WSB mice, WSB.*APP/PS1* mice had no significant changes in PS in individual brain regions or on average.

**Figure 4:**
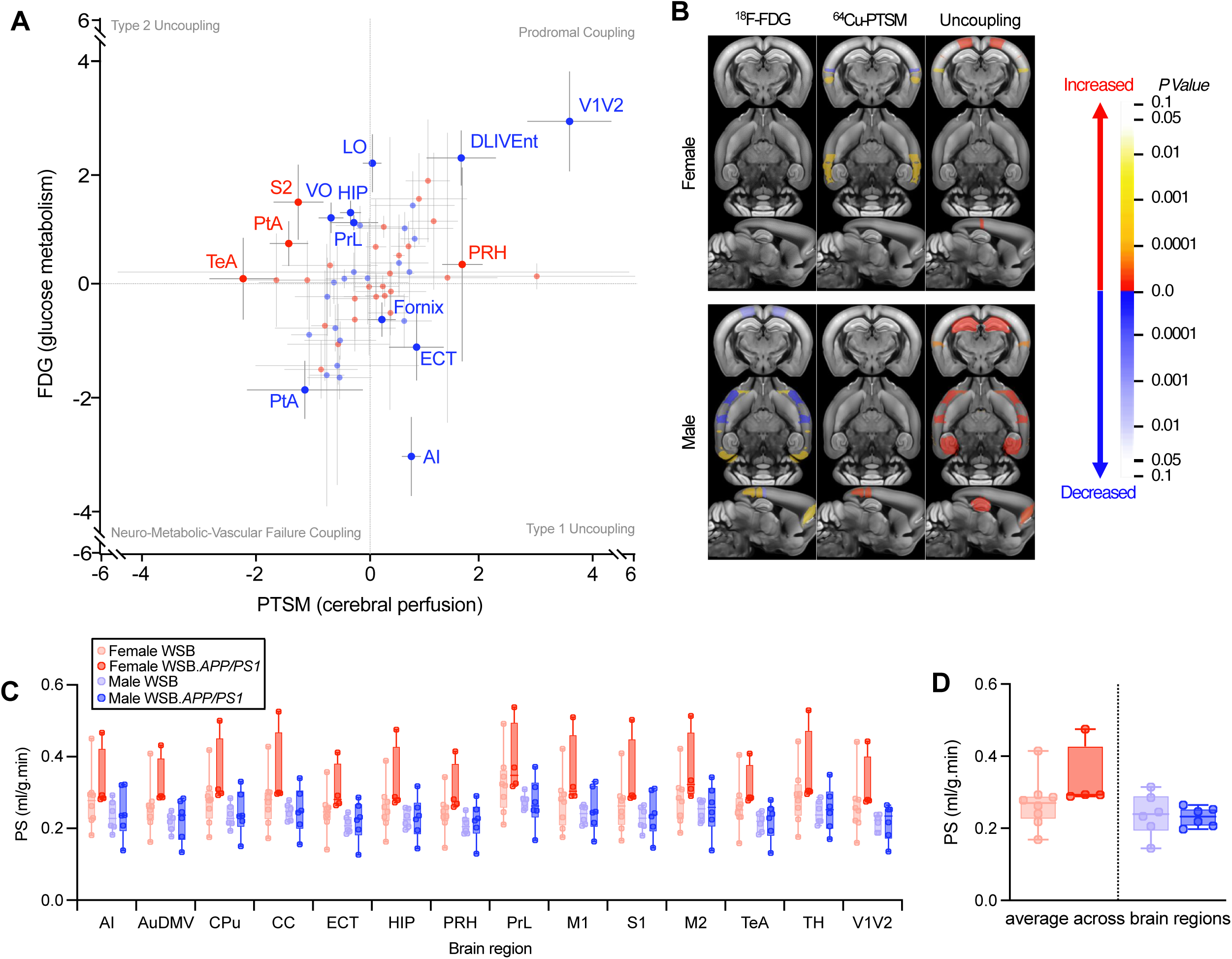
Uncoupling analysis in WSB and WSB.APP/PS1 mice. (**A**) PTSM and FDG uncoupling chart of WSB.*APP/PS1* relative to WSB mice. WSB.*APP/PS1* mice showed regional neurovascular uncoupling (number of regions for female and male, respectively: prodromal: 10, 8; failure: 8, 11; type 1 uncoupling: 7, 4; type 2 uncoupling: 3, 5). Labeled brain regions were significantly different compared to WSB controls, where data are mean±SEM. (**B**) Spatially localized significant regions (*p* < 0.05), where significant regions were projected onto the Allen object maps, with warm colors indicating increases, and cool colors indicating decreases. **(C)** Vascular permeability and surface area (PS) across brain regions in WSB and WSB.*APP/PS1* mice. In both sexes, brain region significantly affected PS (*p* < 0.001 for each; two-way ANOVAs). Genotype did not significantly affect PS for either sex. **(D)** PS averaged across brain regions in WSB and WSB.*APP/PS1* mice. Genotype did not significantly influence average PS for either sex (for females and males: *p* = 0.191 and *p* = 0.806, two-tailed *t* tests). For WT and *APP/PS1*, respectively: female: *n =* 8, 4; male: *n =* 6, 6.

In an effort to better understand changes in vasculature morphology between WSB and WSB.*APP/PS1*, we investigated if total vascular volume and/or exchange surface area changed with genotype and sex through *in vivo* labeling of vessels by DyLight (**Fig. 5A**). Despite no differences in overall brain volume between genotypes (**Fig. 5B**), vascular tree volume was significantly different with genotype for both males and females (**Fig. 5C**). In addition, computing the exchange surface area (S), as the perimeter of the vessel cross-section, revealed that whole brain vascular S was also signficantly higher in WSB.*APP/PS1* mice than WSB (**Fig. 5D**). To understand if this S change was driven by the vessel size and how these were distributed, we computed histograms of all vessel diameters across the entire vascular tree (**Fig. 5E, F**) in a WSB and WSB.*APP/PS1* female mouse. The WSB.*APP/PS1* mouse showed a shift in vessel radii, with 17.7% of vessels with radii between 20.2µm-45.4µm compared to 13.2% in the WSB mouse, consistent with the development of capillaries and meta-aterioles.

**Figure 5:**
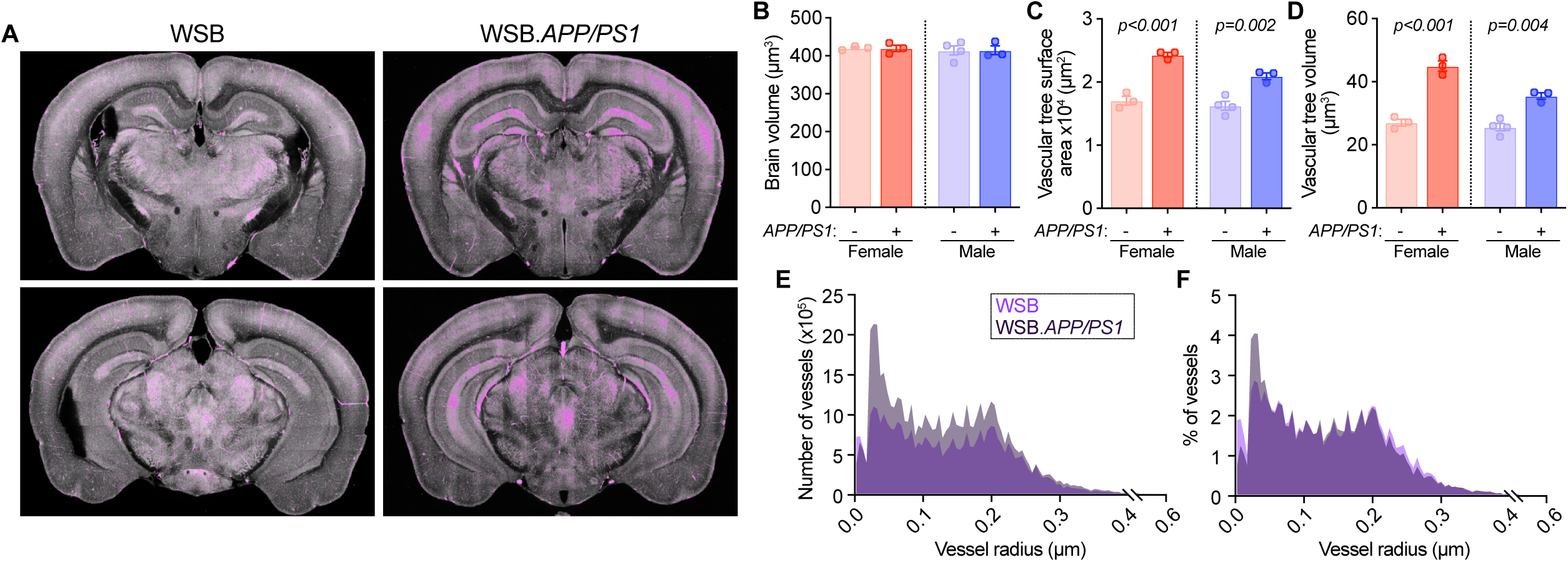
Characterization of extracted vascular tree in WSB and WSB.APP/PS1 mice. (**A**) Representative 2D extracted vascular tree (purple) for WSB and WSB.*APP/PS1.* (**B**) Total brain volume (µm^3^) was not different between WSB and WSB.*APP/PS1* mice for females or males (*p* = 0.953 and *p* = 0.980, respectively; two-tailed *t* tests). (**C**) Vascular tree volume (µm^3^) was significantly increased in WSB.*APP/PS1* brains compared to WSB brains for both females and males (*p* < 0.001 and *p* = 0.002, respectively; two-tailed *t* tests). (**D**) Vascular tree surface area was significantly increased in WSB.*APP/PS1* brains compared to WSB brains in both females and males (*p* < 0.001 and *p* = 0.004, respectively; two-tailed *t* tests). **B-D** For WT and *APP/PS1*, respectively: female: *n =* 3, 3; male: *n =* 4, 3. Data are depicted as mean ± SEM. (**E-F**) Histograms depicting numbers **(E)** and percentages **(F)** of vessels at specific measured radii. *n* = 1 female/genotype.

### WSB vessels did not show age-dependent decline in vascular function and integrity

To better understand the functional cerebrovascular differences specific to WSB, we performed ex vivo studies of posterior cerebral arteries from young (6M) and old (23M) WSB female mice, and compared this with age-matched B6. At young ages, there were no differences in the cerebral artery endothelium-dependent vasodilation, measured as the response to acetylcholine, between B6 and WSB mice (**Fig 6A, B**). Cerebral arteries from old B6 mice had significantly impaired endothelium-dependent vasodilation compared with arteries from young B6 (**Fig 6A, B**). In contrast, arteries from old WSB mice were not impaired, with endothelium-dependent vasodilation similar to that of young B6 and young WSB (**Fig 6A, B**). In all groups, the endothelium-dependent vasodilation to acetylcholine was almost entirely blocked by the nitric oxide synthase inhibitor L-NAME (**Fig 6A**), indicating that nitric oxide bioavailability was lower in old B6, but maintained in old WSB. When examining the response to endothelium-independent vasodilator sodium nitroprusside, we found that all groups had similar responses (**Fig. 6C**), demonstrating that smooth muscle cell responsiveness was not different between groups. The amount of artery preconstriction and maximal artery diameter were similar between groups (**Table 1**), indicating that these potential confounding factors did not impact the vasoreactivity measures. We also measured the ability of the posterior cerebral artery to vasoconstrict in response to endothelin-1 (**Fig 6D, E**). We found that endothelin-1 vasoconstriction was not different between young B6 and young WSB mice. Arteries from old B6 mice vasoconstricted less to endothelin-1 than young B6 arteries. In contrast, the endothelin-1 vasoconstriction was not different between young and old WSB mice (**Fig 6D, E**). Thus, old WSB mice do not have the age-related declines in cerebral artery endothelial function and vasoconstrictor responsiveness that are present in old B6 mice.

**Figure 6:**
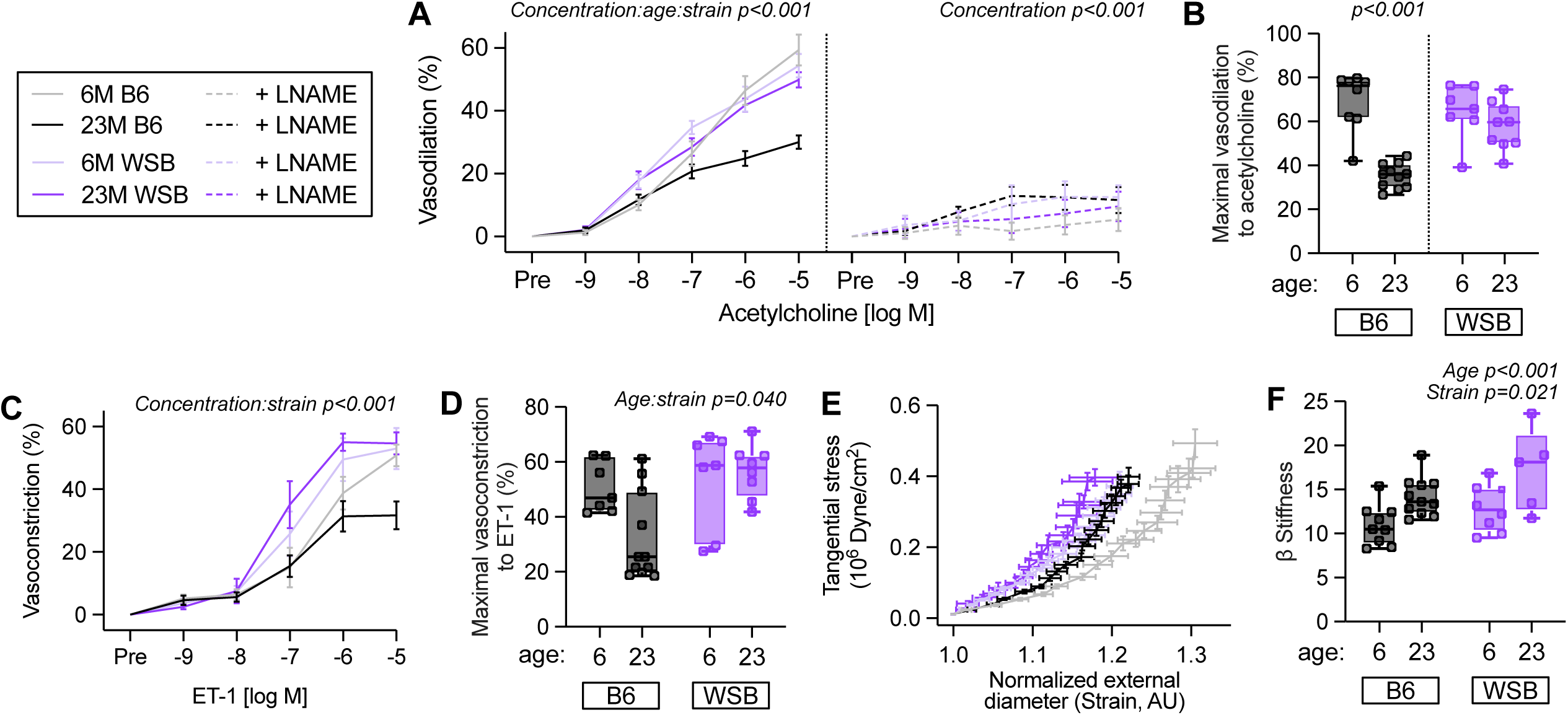
WSB mice did not have age-related impairments in cerebral artery vasoreactivity but did have age-related elevations in cerebral artery stiffness. Isolated posterior cerebral arteries were studied from 6M and 23M B6 and WSB female mice. The vasodilation to endothelium-dependent dilator acetylcholine, displayed as (**A**) dose-response and (**B**) maximal vasodilation, was measured. There was a significant interaction among acetylcholine concentration, strain, and age (*p <* 0.001, three-way ANOVA), but neither strain nor age modified the effect of acetylcholine concentration with the addition of L-NAME (concentration effect: *p <* 0.001, three-way ANOVA). B6 mice had a significantly reduced maximal dilation response with age (*p <* 0.001, Mann Whitney test), but WSB mice did not (*p =* 0.307, two-tailed *t* test; note: strains were analyzed separately due to non-normal distribution). For 6M and 23M, respectively: B6: *n =* 8, 11; WSB: *n =* 7, 9. (**C**) Dose-response to sodium nitroprusside. Neither strain nor age influenced the effect of sodium nitroprusside concentration on vasodilation (concentration: *p* < 0.001, three-way ANOVA). For 6M and 23M, respectively: B6: *n =* 9, 11; WSB: *n =* 10, 9. **(D)** Dose-response and (**E**) maximal vasoconstriction to endothelin-1 (ET-1). Strain significantly modified the effect of endothelin concentration on vasoconstriction (Strain:concentration effect: *p <* 0.001, three-way ANOVA). Strain significantly modified maximal vasoconstriction to ET-1 with age (age:strain effect: *p =* 0.040, two-way ANOVA, Holm-Sidak’s multiple comparison test). B6 mice had significantly decreased maximal vasoconstriction in response to ET-1 with age (*p =* 0.030), which was not observed in WSB mice (*p =* 0.900). Aged WSB mice had significantly higher maximal vasoconstriction compared to aged B6 mice (*p =* 0.003). For 6M and 23M, respectively: B6: *n =* 7, 11; WSB: *n =* 7, 8. (**F**) Stress-strain curve used to calculate β stiffness (**G**). Both age and strain significantly modified β stiffness (Age effect: *p <* 0.001, strain effect: *p =* 0.021 respectively, two-way ANOVA), but strain did not modify age-induced effects on β stiffness (Age:Strain effect; *p =* 0.5988, two-way ANOVA, Holm Sidak’s multiple comparison test). Both B6 (*p =* 0.0499) and WSB (*p =* 0.0456) had increased β stiffness with age. For 6M and 23M, respectively: B6: *n =* 9, 11; WSB: *n =* 8, 5.

**Table 1.** Posterior cerebral artery characteristics for ex vivo studies.

| Variable | Young B6 | Old B6 | Young WSB | Old WSB |
| --- | --- | --- | --- | --- |
| Maximal Diameter ($\mu\text{m}$) | $148 \pm 1$ | $150 \pm 4$ | $135 \pm 6$ | $144 \pm 5$ |
| Maximal SNP dilation (%) | $95 \pm 2$ | $83 \pm 4$ | $90 \pm 3$ | $91 \pm 5$ |
| Preconstriction (%) |  |  |  |  |
| ACh | $24 \pm 2$ | $22 \pm 2$ | $21 \pm 1$ | $18 \pm 2$ |
| ACh + LNAME | $42 \pm 5$ | $37 \pm 4$ | $28 \pm 4$ | $30 \pm 1$ |
| SNP | $36 \pm 5$ | $27 \pm 1$ | $35 \pm 4$ | $33 \pm 3$ |

Data are mean ± SEM. ACh, acetylcholine; SNP, sodium nitroprusside. No statistical differences between groups (all $p > 0.05$ ).

We measured the passive stiffness of posterior cerebral arteries incubated in a solution free from calcium to remove any myogenic tone. We used the changes in lumen diameter and wall thickness in response to increasing pressure to create stress-strain curves (**Fig 6F**) and calculate β-stiffness. We found that old age led to stiffer cerebral arteries in both B6 and WSB mice compared with young mice (**Fig 6G**). This analysis also revealed that WSB mice had stiffer cerebral arteries than B6 mice regardless of genotype (**Fig 6G**). Thus, old age and the WSB strain are associated with stiffer cerebral arteries. Unlike the findings for vasoreactivity, the old WSB mice are not protected against the age-related increases in cerebral artery stiffness.

### Humanized APOE alleles differentially modified CAA, plaque deposition, and plaque-associated microglial area in WSB.APP/PS1 mice

The ε4 allele of *APOE* (*APOE^4^*) is the most prominent risk allele for the development of late-onset Alzheimer’s disease, with the common ε3 allele (*APOE^3^*) being a neutral allele, and ε2 allele (*APOE^2^*) being protective. *APOE^4^* is also known to increase severity of CAA in humans and mouse models(*67–72*). Therefore, to determine the effect of *APOE* status in the context of WSB-specific vascular deficits, a humanized *APOE* allelic series was backcrossed onto the WSB genetic background, generating WSB.*APOE^2/2^APP/PS1,* WSB.*APOE^3/3^APP/PS1*, and WSB.*APOE^4/4^APP/PS1* strains. Male and female mice of each strain were aged to 8M, and brain sections were assessed for CAA and parenchymal plaques, as well as plaque-associated microglia as described previously (**Fig. 1**). Compared to WSB.*APOE^2/2^APP/PS1*, WSB.*APOE^4/4^APP/PS1* had significantly increased CAA regardless of sex, whereas WSB.*APOE^3/3^APP/PS1* had significantly increased CAA in males, but not in females (**Fig 7A-C**). Observationally, using data presented in Figure 1, the degree of CAA in WSB.*APP/PS1* mice (mouse *Apoe; Apoe^WT/WT^*) appeared equivalent to WSB.*APP/PS1* mice carrying either *APOE^2^* or *APOE^3^* (**Fig. 7C**).

**Figure 7:**
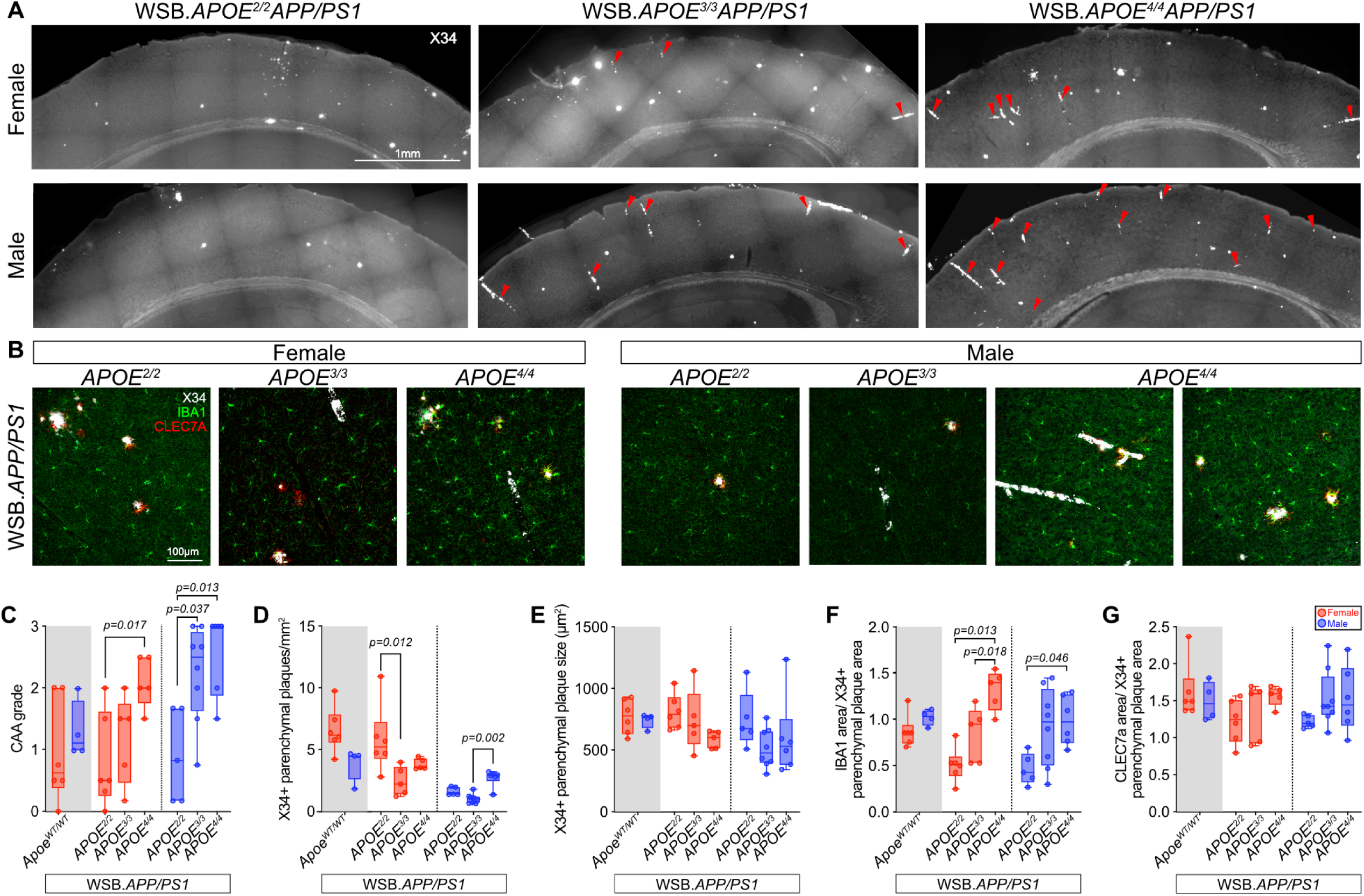
CAA and parenchymal plaque characterization in WSB.APP/PS1 mice expressing APOE ε2, ε3, or ε4 *alleles*. **(A)** Female and male 8M WSB.*APOE^2/2^APP/PS1*, WSB.*APOE^3/3^APP/PS1,* and WSB. *APOE^4/4^APP/PS1* brain hemisections stained with X34 (white) and immunoassayed for IBA1(green) and CLEC7A (red). Arrowheads indicate CAA. Magnification: 20X; scale bar: 1mm. **(B)** High-resolution images of representative parenchymal plaques and CAA. Magnification: 40X; scale bar: 100µm. **(C)** Quantification of CAA scores. *APOE* genotype significantly influenced CAA in both females (*p =* 0.020, one-way ANOVA, Tukey’s multiple comparisons test) and males (*p =* 0.005, Brown-Forsythe ANOVA test, Dunnett’s T3 multiple comparisons test). **(D)** Cortical parenchymal X34+ plaque quantifications normalized to cortical area. *APOE* genotype significantly influenced plaque number in both females and males (*p =* 0.008 and *p <* 0.001 respectively, Kruskal-Wallis tests with Dunn’s multiple comparisons tests). **(E)** Average plaque size quantifications. No differences were observed in either sex across ages (female: *p =* 0.113, one-way ANOVA; male: *p =* 0.150, Kruskal-Wallis test). **(F-G)** Average IBA1+ **(F)** and CLEC7A+ **(G)** area normalized to average plaque size. *APOE* genotype significantly influenced IBA1+ area in both females and males (*p <* 0.001 and *p =* 0.039 respectively, one-way ANOVAs, Tukey’s multiple comparisons tests). No differences were observed for CLEC7A+ area (female: *p =* 0.197, one-way ANOVA; male: *p =* 0.514, Brown-Forsythe ANOVA test). For WSB.*APOE^2/2^APP/PS1*, WSB.*APOE^3/3^APP/PS1*, and WSB.*APOE^4/4^APP/PS1*, respectively; female: *n =* 6, 5, 5; male: *n =* 5, 8, 6. Data highlighted in gray are recapitulated from respective 8M WSB.*APP/PS1* measurements reported in Figure 1 (all of which had WT mouse *Apoe* genotypes; *Apoe^WT/WT^*) and were not statistically analyzed alongside WSB.*APOE*/* APP/PS1* data, as they were generated separately.

*APOE* genotype also significantly modified parenchymal plaque deposition in both males and females (**Fig. 7D**). In females, *APOE^3^*reduced parenchymal plaque deposition compared to *APOE^2^*, and in males, *APOE^4^* had more parenchymal plaques compared to *APOE^3^*. *APOE* genotype did not significantly modify parenchymal plaque size in males or females (**Fig. 7E**). To investigate whether humanized *APOE* modulated plaque-associated microglial volume, IBA1+ microglia and CLEC7A+ area was measured within a 30µm radius from the edge of X34+ parenchymal plaques. Interestingly, *APOE^4^* genotype significantly increased plaque-associated microglial volume compared to *APOE^2^*in both males and females (**Fig. 7F**). *APOE^3^* genotype also significantly increased plaque-associated microglial volume compared to *APOE^2^* in females (**Fig. 7F**). Plaque-associated CLEC7A+ area was consistent across *APOE* genotypes regardless of sex (**Fig. 7G**). Therefore, *APOE^4^*resulted in both increased CAA and parenchymal plaque-associated microglial volume.

## DISCUSSION

AD is considered the most common form of dementia, and vascular anomalies, such as CAA, are present in most cases. Despite this, many mouse models relevant to AD have focused on the primary AD pathologies—amyloid and tau. For instance, the commonly used transgenic-based 5xFAD(*73*) and *APP/PS1* models, as well as the recently developed knock-in *APP^SAA^*(*41*) and NL-G-F(*74*) models show small but significant amounts of CAA, primarily at later ages (*41*). To date, all of this work has been done on the C57BL/6 background. In mice, CAA has been studied using rare mutations that model CAA, such as the Dutch or Iowa mutations(*75*), although the pathophysiology in these cases are unlike clinical presentation seen in AD. The lack of co-occurrence between hallmark AD pathologies (amyloid and tau), and vascular anomalies (e.g., CAA) in commonly used mouse models has likely hampered our abilities to understand how they synergize to cause neurodegeneration and cognitive decline. In this study, we continue to show the value of non-traditional inbred mouse strains to study AD. We previously reported that WSB.*APP/PS1* showed significant levels of CAA and cognitive deficits at 8 months(*9*), that correlated with differences in myeloid cell profiles(*10*). Here, we define the timeline of CAA development and show they have human AD-relevant neurovascular and metabolic uncoupling. We also use transcriptomics to define pathways perturbed throughout the disease course, including myelin deficits, and confirm this using immunofluorescence. Finally, we create an allelic series of humanized *APOE* in combination with *APP/PS1* on the WSB background and show that *APOE4* increased the degree of CAA.

Multiple studies suggest changes to energy metabolism(*76–78*) and vascular health(*63, 79*), particularly in cerebral blood flow, occur prior to any signs of cognitive decline – suggesting these may be early drivers of AD. Similarly, PET/CT from WSB.*APP/PS1* mice showed vascular and metabolic uncoupling occurs early in the disease course. Clinical studies have demonstrated a reduction in glucose brain metabolism and cerebral blood flow disturbances in at-risk patient populations even before detectable levels of amyloid accumulation(*80–82*). Initially, blood flow may be increased in response to an energy deficit, as has been observed both preclinically and clinically. In response to blood flow disturbances, a strong angiogenic response can occur to counter the effects of low oxygen tensions, leading to an increase in vascular density(*83*). Arteriogenesis can also occur as it is driven by hemodynamic factors such as stretch and shear stress(*84*). Disordered vascular remodeling and arteriovenous malformations can also lead to blood vessel rupture and organ hemorrhage. Interestingly, studies in stroke(*85, 86*) and traumatic brain injury(*87*) have shown that cerebral perfusion insufficiency results in a metabolic depression and concomitant hypoperfusion. Central to our hypothesis is that age-dependent reductions in cerebral blood flow lead to regional decreases in glycolytic metabolism.

Until recently, CAA has been largely absent in the majority of AD models. In WSB.*APP/PS1,* CAA first appears between 4 and 8 months, with more significant CAA apparent by 14 months, which coincides with the emergence and progression of parenchymal amyloid. To quantify CAA, we modified a method developed to grade CAA in human postmortem brains that was based on clinical scoring methods. The presence of CAA in the leptomeninges received the lowest score (score 0.5) with CAA in the penetrating cortical vessels increasing the CAA score based on the number of vessels affected (score 1-3). Although not quantified, the presence of CAA in WSB.*APP/PS1* showed regional variability across mice, with different cortical regions more or less affected. This aligns with human studies that show AD patients can show varying degrees of CAA, with certain cortical regions (e.g., occipital and parietal cortex) often most affected(*88*). Despite its effectiveness at relatively quickly and efficiently ‘grading’ CAA, improved quantification methods for CAA are needed, and machine-learning approaches are under devleopment by our group, with the intent to uncover the mechanistic drivers of CAA in WSB.*APP/PS1*. We anticipate the WSB strain harbors genetic factors that increase risk for CAA and possibly other vascular anomalies. Genetic mapping approaches – such as quantitative trait loci mapping – can facilitate the identification of these factors. Interestingly, previous studies showed that genetic loci in the WSB strain modify outcome measures in a stroke model (*89*). Studies are underway using a B6/WSB F2 mapping population to identify the genetic factors that modify risk for CAA, and it will be interesting to compare with those identified in the stroke model. Ultimately, these genetic factors will inform us about the mechanisms driving CAA in AD, and hopefully identify novel therapeutic avenues to explore to treat AD.

To begin to understand the mechanisms underlying vascular dysfunction in the WSB context, we tested age-related vessel function, contrasting young and old WSB with B6 mice. Our findings for lower endothelium-dependent vasodilation and endothelin-1-mediated vasoconstriction in old B6 mice are consistent with the literature(*44, 90–93*). Consistent with previous studies, we also find that the impairment in endothelium-dependent vasodilation is due to reduced nitric oxide bioavailability(*44, 91, 92*). Previous studies have further demonstrated that increased reactive oxygen species are the primary driver of the lower nitric oxide bioavailability in old age(*45, 91*). Endothelin-1 also increases in the plasma and endothelial cells with age (*94*). It has been proposed that these higher levels of endothelin-1 cause vasoconstriction impairments due to desensitization(*95*). Importantly, these studies demonstrate that old WSB mice do not have the same impairments in vasodilation and vasoconstriction as old B6 mice. These are potentially mediated by lower oxidative stress and endothelin-1 in old WSB, which requires further investigation. Unlike the vasoreactivity measures, WSB cerebral arteries are vulnerable to age-related stiffening. The functional consequences of cerebral artery stiffening are not entirely clear. It is known that the age-related stiffening of large extracranial arteries is associated with cerebral artery dysfunction(*96, 97*), likely due to the resulting elevation in pulse pressure. However, it is unknown if the stiffness of the cerebral arteries affects their own vasoreactivity. Cerebral arteriole stiffness is also related to impaired glymphatic function(*98*), a potential pathway for Aβ clearance from the brain(*99*). Thus, the greater cerebral artery stiffness in the WSB mice could impact Aβ clearance, an area warranting further study.

Vascular deficits have only more recently been appreciated as critical contributors to many dementias, including AD, mixed (etiology) dementia, Parkinson’s disease and others. Vascular deficits include a variety of pathologies—including CAA, blood flow deficits, lacunar infarcts and small vessel disease (SVD). Mechanisms underlying vascular contributions to cognitive decline and dementia (VCID) are not well understood, and this may be contributing to the lack of success in clinical trials for particularly AD. Few if any clinical trials are focusing on vascular pathologies (*100*). In addition, current anti-amyloid-based treatments increase risk for amyloid-related imaging abnormalities (ARIA), that are thought to be because of cerebrovascular deficits(*101*). APOE4 significantly increases risk of ARIA, possibly due to the increased CAA(*101*). Given the need to develop strategies to treat VCID and ARIA, improved mouse models that recapitulate these key features are essential. We propose incorporation of WSB genetic context into mouse models will improve the translatability of the next generation of ADRD mouse models. To this end, WSB.*APP/PS1* are available from the JAX repository (#33567283) and the *APOE* allelic series on WSB are available upon request to the corresponding author. In addition, the IU/JAX/PITT MODEL-AD (Model Organism Development and Evaluation for Late-onset AD) center is including WSB as a second genetic context in its collection of LOAD mouse models. MODEL-AD is charged with creating preclinical mouse models to improve the translatability of preclinical studies to the clinic(*102*). Models incorporate aging as well as genetic and environmental risk factors to recapitulate the complexity of human LOAD. Although models are standardized on the B6 background, a subset of those models is being generated on the WSB background, incorporating *APOE4* in combination with WT and mutant humanized Aβ and *MAPT* alleles. All models, as well as all associated phenotyping data, are made available at the earliest opportunity.

In summary, we show that WSB.*APP/PS1* mice developed age-related parenchymal plaque deposition and CAA, which was modulated by *APOE* genotype. Transcriptomic analysis indicated that WSB.*APP/PS1* had significant overlap to changes observed in human AD, highlighting the relevance of WSB.*APP/PS1* mice. Further, WSB.*APP/PS1* mice exhibited a loss of myelin, metabolic uncoupling, increased vascular tree volume, with the WSB background having increased artery stiffness, and differential resilience to age-related decline in vascular responsiveness compared to B6—each of which may be important facets of cerebrovascular deficits in AD warranting further study.

## AUTHOR CONTRIBUTIONS

Olivia J. Marola, Gareth R. Howell, and Kristen D. Onos conceived of the idea for the manuscript. Olivia J. Marola, Ashley E. Walker, Paul R. Territo, Gareth R. Howell, and Kristen D. Onos designed experiments. Olivia J. Marola, Asli Uyar, Kelly J. Keezer, Kevin J. Elk, Jonathan Nyandu Kanyinda, Juan Antonio K. Chong Chie, Scott Persohn, Paul R. Territo, Paul Salama, Jennifer D. Whitesell, Julie A. Harris, Ashley E. Walker, Abigail E. Cullen, and Kristen D. Onos performed experiments. Olivia J. Marola, Asli Uyar, Juan Antonio K. Chong Chie, Ashley E. Walker, Gregory W. Carter, Michael Sasner, Paul R. Territo, Gareth R. Howell, and Kristen D. Onos analyzed and interpreted data. Olivia J. Marola, Asli Uyar, Juan Antonio K. Chong Chie, Ashley E. Walker, Paul R. Territo, Gareth R. Howell, and Kristen D. Onos drafted the manuscript. All authors edited and approved the manuscript.

## ACKNOWLEDGEMENTS

We are grateful to JAX Scientific services, particularly Genome Technologies, Transgenic Genotyping Services and Jarek Trapszo from Scientific Instrumentation Services. We acknowledge Phillip Bohn for his contributions to vascular 3D labeling and imaging. This work was funded by U54-AG054345 (GRH, PRT, GWC), U54-AG054349 (GRH, PRT, GWC), R21-AG078575-01 (GRH, PRT), R01-AG064016 (AEW), R01-AG047589 (JAH), and AARF-22-971325 (OJM). GRH holds the Diana Davis Spencer Foundation Chair for Glaucoma Research and GWC holds the Bernard and Lusia Milch Endowed Chair.

## CONFLICT OF INTEREST STATEMENT

All authors declare that they have no conflicts of interest to disclose.

## CONSENT STATEMENT

No human subjects were used in the current study, and therefore consent was not necessary.

## DATA AVAILABILITY STATEMENT

RNA-sequencing data are available via the AD Knowledge Portal with Synapse ID #####. All other data are available from the corresponding authors upon reasonable request.

---

**Figure S1:**
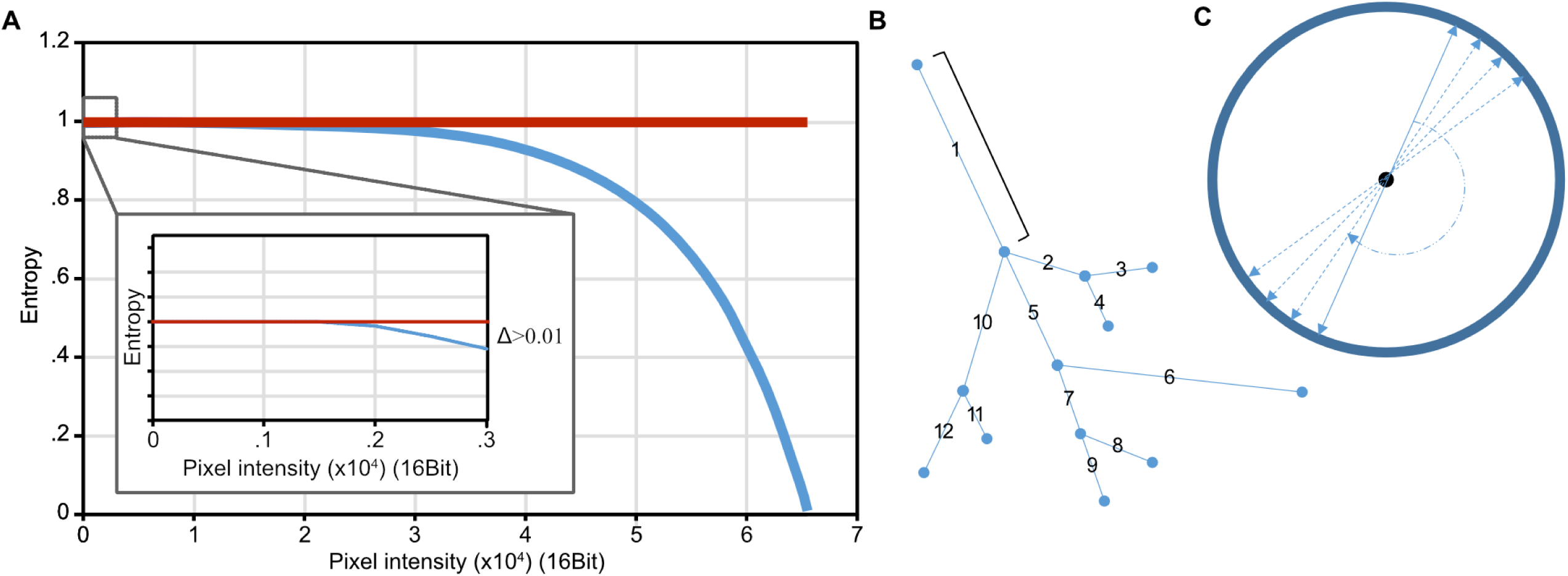
3D Vessel Labeling, Imaging, and Quantification Methods. **(A)** Example graph of entropy cutoff. Data plotted as entropy vs. intensity range (spanning from 0 to 65535) were used to determine the point where the entropy curve starts to decrease (i.e. Δ>0.01), which was selected as the minimum of the threshold range and used to obtain a coarse initial segmentation of the vessels’ volume. **(B)** Schematic of branch counting of the skeletonized vascular tree depicting locations where the vessels split; branches are defined as a segment, or connected voxels, between two splitting locations. **(C)** Ray tracing for diameter and perimeters. A ray casting process is used to calculate the vascular exchange surface (perimeter of the slice) and cross-sectional areas, where origin of the ray was centered along the midline, and the ray in each direction intersects the vessel contour 180 out of phase. This process is repeated iteratively by rotating the angle between rays over the interval of 0° to 180° with 1° steps. The algorithm returns a set of measurements for each voxel in the skeletonized vessel tree.

## Notes

### Competing Interest Statement

The authors have declared no competing interest.

### Summary of Updates

This version has updated figures that were difficult to view, and clarified the text.

